# Awareness is determined by emotion and gender

**DOI:** 10.1101/2024.02.27.579317

**Authors:** Ema Jugović, Marta Poyo Solanas, Beatrice de Gelder

**Affiliations:** Department of Cognitive Neuroscience, Facult y of Psychology and Neuroscience, Maastricht University, Maastricht, The Netherlands

**Keywords:** awareness, bodies, emotion, features, gender, masking

## Abstract

Traditionally, consciousness studies focus on domain general cognitive processes rather than on specific information reaching subjective awareness. The present study (*N* = 45) used visual masking and whole-body images to investigate whether the specific emotional expression as well as gender of the stimuli and of the participants impact awareness. Our results show that participants’ awareness responses reflect differences in the specific emotion of the stimuli, that these differences are a function of the gender of the stimuli as well as the gender of the participants and that minimal awareness may be associated with emotion specific features of the body images. Overall, we observed that threatening expressions are more easily detected than fearful ones, especially by males presented with male stimuli. Our findings underscore the importance of affective factors for theories of consciousness and underscore the significance of gender differences in emotional processing, often overlooked in past face and body emotion recognition studies.

## 1 Introduction

Survival in rich and complex environments requires rapid detection of unexpected events, such as social threat, and builds on the ability to react adaptively to them. It is currently a matter of debate whether adaptive reactions to social threat require full conscious perception of the stimuli. Current theories of consciousness predominantly address cognitive determinants of awareness at the detriment of affective factors. This leaves little room for understanding whether affective factors also play a role. Yet, affective stimuli may be prioritized in accessing awareness because of their behavioral relevance (Tamietto and de Gelder, 2008; Tamietto and de Gelder, 2010; Striemer, Whitwell and Goodale, 2013; Tamietto et al., 2015). For instance, several studies reported fearful faces entering visual awareness faster than other expressions (Yang, Zald and Blake, 2007; Gray et al., 2013; Stein et al., 2014). Using continuous flash suppression (CFS), shorter suppression times were revealed for fearful facial expressions compared to happy and angry (Gray et al., 2013), happy and neutral (Yang et al., 2007) and neutral faces (Stein et al., 2014). Esteves and Öhman (1993) also found evidence of happy facial expressions having perceptual precedence over angry and neutral faces using a visual backward masking paradigm. Findings about affective determinants of awareness are more compatible with theories that trace consciousness to biological and somatic processes (Panksepp, 2007; Damasio, 1999). On the strength of such theories one expects that subjective awareness is selectively sensitive to different emotional signals and perceivers.

Similarly to facial expressions, emotional whole-body expressions represent an equally salient social and affective signal, yet they have not received as much attention in studies involving perceptual awareness of affective stimuli. But important differences in non-conscious processing of faces and bodies have been found (Zhan, Hortensius and de Gelder, 2015). Using CFS, Zhan et al. (2015) demonstrated that angry facial expressions are suppressed from conscious perception longer than fearful or neutral faces, whereas the effect was reversed for bodies. Angry body expressions broke from suppression faster than fearful and neutral bodies (Zhan et al., 2015). Interestingly, emotional body postures were suppressed from awareness longer than emotional facial expressions. These findings concerning body expressions have been further replicated in a recent study by Poyo Solanas, Zhan and de Gelder (2023) employing a similar paradigm. Taken together, these results illustrate the importance of investigating different sources of emotional information.

Another issue is that gender differences might play a significant role in affective visual awareness. Several behavioural studies investigating emotion recognition from faces and bodies reported significant gender effects (Sokolov et al., 2011; Lambrecht et al., 2014). For instance, Lambrecht et al. (2014) found that females exhibit greater recognition of happy, alluring and neutral facial expressions, while a similar non-significant trend was observed for disgust and anger. Moreover, multiple studies demonstrated higher accuracy in recognizing emotional prosody for females (Lambrecht et al., 2014; Lausen and Schacht, 2018). Further, Sokolov et al. (2011) presented happy, angry and neutral door-knocking actions via point-light displays and found greater recognition accuracy of happy and neutral actions in males and females, respectively. Nonetheless, one study failed to find any gender differences in perceiving emotional body motion (Isernia et al., 2020).

Neuroimaging studies also provide evidence for gender differences in affective processing (Schneider et al., 2000; Hofer et al., 2006; Kret et al., 2011; He et al., 2018). In an event-related potentials study, females had significantly larger P1 components when observing male compared to female angry whole-body expressions, whereas males revealed greater P3 amplitudes for female than male angry body postures (He et al., 2018). An fMRI study by Kret et al. (2011) observed increased brain activity for threatening bodies in the superior temporal sulcus (STS), extrastriate body area (EBA) and pre-supplementary motor area (pre-SMA) when male participants observed male in comparison with female actors. Other fMRI studies employing emotion-inducing paradigms reported differential brain region activations among males and females during pleasant and unpleasant emotions (Hofer et al., 2006) and significant amygdala activation for induced sadness in males, but not females (Schneider et al., 2000).

Another issue is whether there is a relation between perceptual awareness and whole vs. part-based perception. Faces and bodies are presumably processed configurally rather than by isolated parts or features. In this regard, research has shown that configuration-based processing is disrupted in favor of feature processing when inverting face (Tanaka and Farah, 1993) and body stimuli (Reed et al., 2003; Bannerman et al., 2009; Stekelenburg and de Gelder, 2004), leading to poorer recognition performance. Similarly, emotion recognition from faces and whole-body expressions is impaired with stimulus inversion (Balas and Huynh, 2015; Stein, Sterzer and Peelen, 2012). Stein et al. (2012) also found significantly longer breaking from suppression times for inverted compared to upright human faces and bodies using the CFS paradigm. This delay suggests that stimulus inversion interferes with access to consciousness (Stein et al., 2012). Since inversion disadvantages configural processing in favor of part-based processing, this pattern may indicate configural processing in the early stages of awareness. On the other hand, processing outside or with minimal awareness may be driven by feature rather than configural processes. Body expressions differ from each other in specific features as much as in overall configuration. For example, limb contraction contributes significantly to fearful body expression perception (Poyo Solanas, Vaessen and de Gelder, 2020a). Likewise, postural features, such as symmetry, also correlate with other specific emotional categories (Poyo Solanas et al., 2020a,b). Perception of some critical features may be sufficient to trigger early expression specific perception under conditions of limited visibility, at least as much as is needed for triggering adaptive behavior (de Gelder and Poyo Solanas, 2021). If so, this may provide evidence for feature-based discrimination between the expressions in early visibility stages and is a relevant aspect of minimal awareness.

Against this background, the present study addresses three outstanding questions. Firstly, do different emotional whole-body expressions trigger differences in awareness? Secondly, does the gender of the stimuli and that of the participants also play a role? Finally, how do the results support configural or feature-based perception in minimal awareness?

## 2 Materials and methods

### 2.1 Participants

The sample consisted of 45 participants (22 males, mean age = 23.8 ± 6.35 years, age range = 18 – 48 years). All participants had normal or corrected-to-normal vision, were screened for hand dominance and had no medical history of psychiatric or neurological disorders. Study participation was voluntary and carried out with the understanding and written consent of each participant. All procedures followed the regulations of the Ethical Committee at Maastricht University and were conducted in compliance with the Declaration of Helsinki. Participants received credit points for their participation.

### 2.2 Stimuli

A previously validated stimulus set consisting of full-body images with the faces blurred (Stienen and de Gelder, 2011) was adapted for this study using Adobe Photoshop (Adobe Inc., 2023, Adobe Photoshop, ver. 24.7.0). The smallest possible rectangle that fitted all body stimuli was defined, after which all stimuli were centered and adjusted to a similar size. Forty-eight grayscale images of whole-body expressions were used. The body postures portrayed three different emotions, i.e., threat, fear and sadness. Each emotional category consisted of 16 stimuli of the same 16 actor identities (eight males, eight females). All stimuli subtended approximately 5° × 10° of visual angle (210 × 456 pixels) and had a mean luminance of 113.80 ± 6.13. For each stimulus, a corresponding mask was constructed using custom MATLAB scripts (Borgomaneri et al., 2023). This was achieved by randomly redistributing pixels within the original stimulus image, with the objective of preserving low-level visual features, such as spatial frequency and contrast. In addition, 180° inverted versions of each stimulus were created to investigate configural vs. featural bases of minimal awareness.

### 2.3 Task and procedure

The experiment used a visual masking task, adapted from Borgomaneri et al. (2023). The experimental session consisted of 2 runs including 10 practice trials at the beginning of every run. Each run contained of 96 trials divided in 3 blocks (16 stimulus identities x 2 orientations x 3 emotions). All stimuli were presented in a random order.

Each trial started with a baseline period during which a grey screen was shown for ∼800ms, followed by a body image centrally presented for 8ms and immediately followed (SOA = 0) by a mask shown for 16ms (see *Stimuli* for details). Subsequently, participants were instructed to indicate whether they had seen a body or not by pressing “J” for ‘Seen’ and “K” for ‘Unseen’ using their dominant hand. Importantly, participants were instructed to only press ‘Seen’ when they were sure that they had seen a whole-body outline and not just a body part e.g., only the legs or the arms. Instructions emphasized speed as well as accuracy. As soon as the participant responded, the experiment proceeded to the next trial.

To further reduce awareness, white noise was overlaid on the body stimulus and the mask using the “imnoise” MATLAB function (parameter specifications: ‘Gaussian’, mean = 0, starting variance value = 0.1). Importantly, to tackle substantial individual differences regarding stimulus visibility, a staircase procedure was implemented as follows. After the first 10 trials of the task, the amount of white noise on a given trial was determined by the percentage of ‘Seen’ trials in the last 10 trials of the task. If the number of ‘Seen’ trials in the previous 10 trials was smaller or larger than 50 %, the amount of white noise would decrease or increase in increments of 0.2 in the following trial, respectively. This procedure was implemented to obtain approximately equal numbers of seen and unseen trials across participants.

The task was presented in MATLAB R2021b (MathWorks Inc., 2021) using Psychtoolbox-3 version 3.0.18 (Brainard and Vision, 1997; Pelli and Vision, 1997) on an LCD screen (LG Philips LP173WF1, resolution = 1920 × 1080 pixels, screen width = 40.64 cm, screen height = 22.8 cm, refresh rate = 120 Hz) while participants were seated comfortably in front of a screen, approximately 55 cm away.

### 2.4 Data analysis

Data analysis was performed in SPSS (IBM, ver. 28.0, 2021). Outcome measures of interest were stimulus Visibility and Reaction time (in milliseconds). Reaction time outliers were excluded within participants based on the mean ± 2.5 SD criterion.

Visibility and Reaction times were analyzed using Generalized Estimating Equations (GEEs). Due to data averaging across trials of the same condition, the binary Visibility measure took on decimal values within the 0 to 1 interval (mean = 0.422 ± 0.270). Therefore, the Visibility data was modelled with a Gaussian distribution with an identity link function. Conversely, Reaction times data were modelled with a gamma distribution combined with a log link function. Both models employed within-subject factors emotion (threat, fear, sadness), stimulus orientation (upright, inverted) and stimulus gender (female, male). The Unstructured (UN) covariance matrix for repeated measurements was used based on the QIC criterion (Pan, 2001). Statistically significant effects were followed up with post hoc pairwise contrasts, while the significance level was Sidak corrected for multiple comparisons.

## 3 Results

### 3.1 Visibility

To investigate whether visibility of whole-body emotional expressions differs across emotional categories, stimulus orientation as well as stimulus and participant gender, a GEEs model was defined. We found significant main effects of emotion (χ^2^(2, *N* = 45) = 180.88, *p* < .001), stimulus orientation (χ^2^(1, *N* = 45) = 114.75, *p* < .001), stimulus gender (χ^2^(1, *N* = 45) = 29.30, *p* < .001) and significant emotion × stimulus orientation (χ^2^(2, *N* = 45) = 86.98, *p* < .001), emotion × stimulus gender × participant gender (χ^2^(2, *N* = 45) = 9.32, *p* = .009) and stimulus orientation × stimulus gender × participant gender (χ^2^(1, *N* = 45) = 5.49, *p* = .019) interactions.

In contrast, effects of participant gender (χ^2^(1, *N* = 45) = 1.19, *p* = .276), emotion × stimulus gender (χ^2^(2, *N* = 45) = 5.19, *p* = .075), emotion × participant gender (χ^2^(2, *N* = 45) = 2.69, *p* = .260), stimulus orientation × stimulus gender (χ^2^(1, *N* = 45) = 3.33, *p* = .068), stimulus orientation × participant gender (χ^2^(1, *N* = 45) = 1.83, *p* = .176) and stimulus gender × participant gender (χ^2^(1, *N* = 45) = 2.35, *p* = .126) were not significant.

Upon inspection of the significant emotion × stimulus orientation interaction (Figure 1), significant differences in stimulus visibility across the three emotions were found. Specifically, upright presentation of the stimuli resulted in significantly higher visibility for sad stimuli (*M* = .67, *SEM* = .03) compared to threatening (*M* = .58, *SEM* = .03, *t(*89) = −10. 78, *p* < .001) and fearful stimuli (*M* = .40, *SEM* = .03, *t(*89) = −13.74, *p* < .001). Likewise, stimulus visibility was higher for threatening compared to fearful stimuli (*t(*89) = 9.26, *p* < .001).

**Figure 1.**
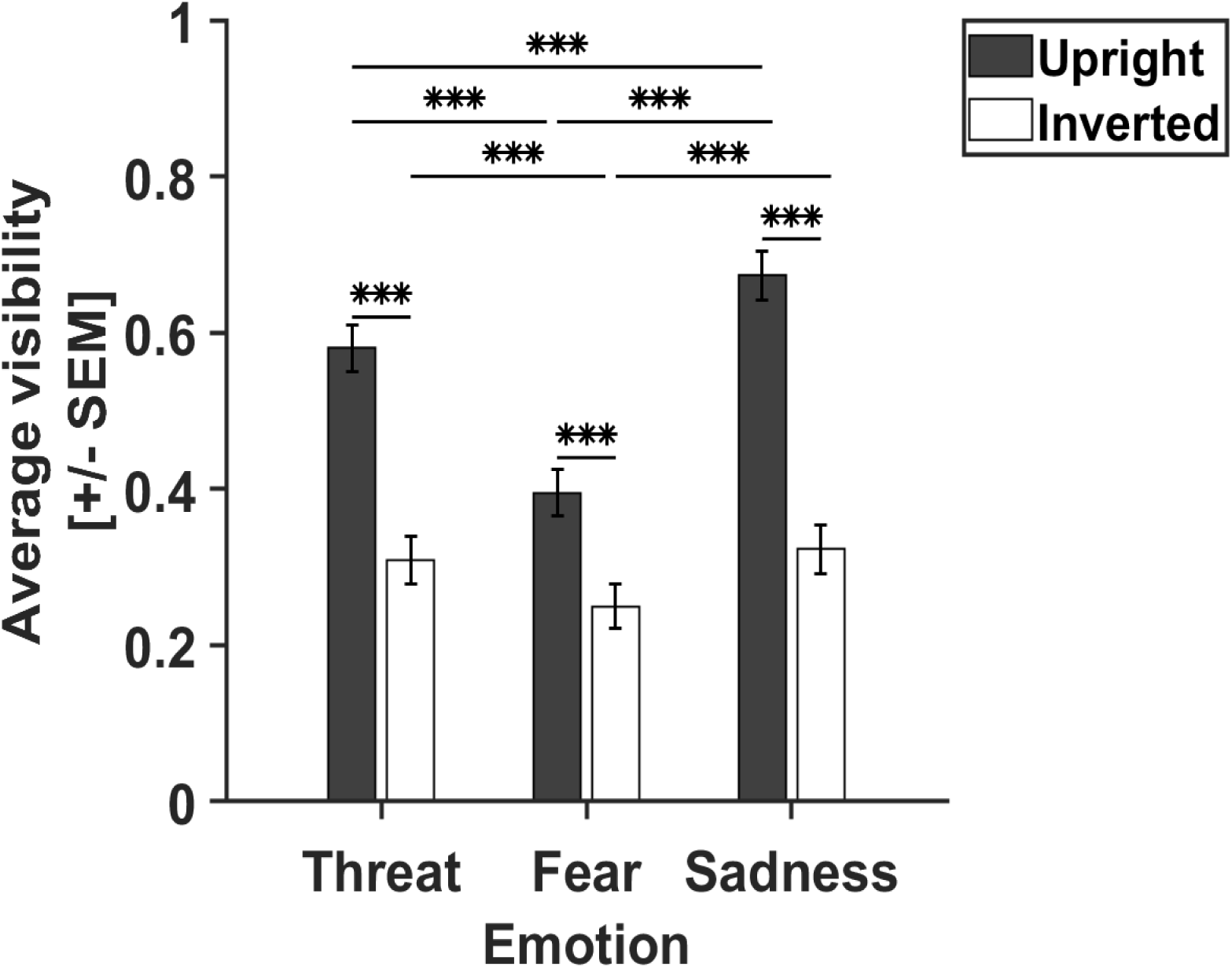
Significant differences in visibility across emotional categories. The x-axis depicts threatening, fearful and sad whole-body expressions, while average stimulus visibility is plotted on the y-axis. Higher values represent higher visibility. Grey and white bars represent upright and inverted stimulus orientation, respectively. Stimulus visibility is highest for sad body postures, followed by threatening and fearful whole-body expressions. Interestingly, visibility across emotion categories is similar for both stimulus orientations, while visibility of the emotional body expressions is significantly reduced with stimulus inversion. *SEM* = standard error of the mean, *** *p* ≤ .001

When stimuli were inverted, a similar trend was maintained (Figure 1). Visibility was significantly higher for sad (*M* = .32, *SEM* = .03) compared to fearful (*M* = .25, *SEM* = .03, *t(*89) = −5.02, *p* < .001), however not compared to threatening whole-body expressions (*M* = .31, *SEM* = .03, *t(*89) = .95, *p* = .719). Likewise, stimulus visibility was significantly higher for threatening (*M* = .31, *SEM* = .03) in comparison with fearful expressions (*t(*89) = 4.16, *p* < .001). Importantly, visibility was significantly reduced following stimulus inversion for all three emotional categories (threat: *t(*89) = 8.98, *p* < .001, fear: *t(*89) = 7.10, *p* < .001, sadness: *t(*89) = 11.92, *p* < .001).

In summary, whole-body expression perception under conditions of reduced visibility depended on both emotional category and upright/inverted stimulus presentation (Figure 1). Despite significantly lower average visibility of inverted stimuli compared to upright, comparable visibility trends across the emotions were observed for both stimulus presentation types. Specifically, sad expressions were significantly more visible than the others in noisy perceptual settings.

Furthermore, follow-up analyses of the significant emotion × stimulus gender × participant gender interaction yielded significant differences in stimulus visibility for the two stimulus gender categories (Figure 2). Namely, male participants reported significantly higher visibility for threatening male (*M* = .55, *SEM* = .04) compared to threatening female stimuli (*M* = .46, *SEM* = .03, *t(*43) = −3.96, *p* < .001) and significantly higher visibility for fearful male (*M* = .47, *SE*M = .04) in comparison with fearful female whole-body expressions (*M* = .40, *SEM* = .03, *t(*43) = - 5.23, *p* < .001). Interestingly, no significant differences between stimulus genders were observed for sad body postures (female stimuli: *M* = .54, *SEM* = .04, male stimuli: *M* = .56, *SEM* = .03, *t(*43) = - 0.93, *p* = .352).

**Figure 2.**
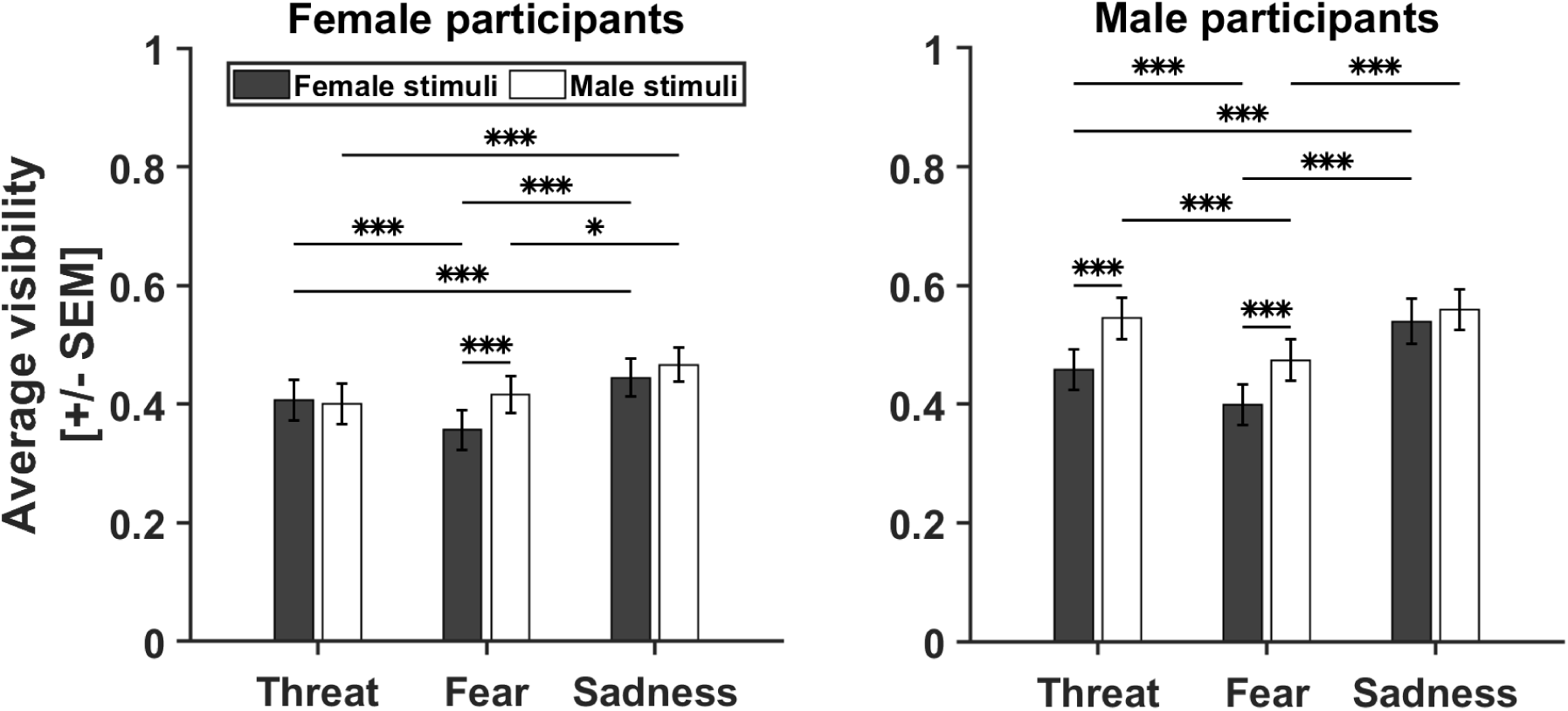
Male participants are more susceptible to male threat. Left and right figure represent data from male and female participants, respectively. The three emotion categories are plotted on the x-axis, while average stimulus visibility is plotted on the y-axis. Higher values represent higher visibility. Grey and white bars depict female and male stimuli, respectively. Male participants reported significantly higher visibility for male threatening and fearful whole-body expressions in comparison with conditions when threatening emotions were expressed by female actors. Conversely, female participants only exhibited significantly higher visibility for fearful male body expressions compared to female. *SEM* = standard error of the mean, *** *p* ≤ .001, * *p* ≤ .05

On the other hand, female participants demonstrated significantly higher visibility only for male fearful (*M* = .42, *SEM* = .03) compared to female whole-body expressions of fear (*M* = .36, *SEM* = .03, *t(*45) = −3.62, *p* < .001, Figure 2). The effect was non-significant for threatening (female stimuli: *M* = .41, *SEM* = .03, male stimuli: *M* = .40, *SEM* = .03, *t(*45) = 0.38, *p* = .702) and sad body postures (female stimuli: *M* = .44, *SEM* = .03, male stimuli: *M* = .47, *SEM* = .03, *t(*45) = - 1.16, *p* > .244) in female participants.

We also inspected visibility differences between emotion categories for the two stimulus genders. For female stimuli, male participants showed significantly higher visibility for female sad (*M* = .54, *SEM* = .04) in contrast with female threatening (*M* = .46, *SEM* = .03, *t(*43) = - 4.57, *p* < .001) and female fearful whole-body expressions (*M* = .40, *SEM* = .03, *t(*43) = −7.50, *p* < .001). Additionally, male participants had higher visibility for female threatening when compared to female fearful body postures (*t(*43) = 4.00, *p* < .001). Similarly, stimulus visibility was significantly greater for male sad (*M* = .56, *SEM* = .04) than for male fearful body expressions (*M* = .47, *SEM* = .04, *t*(43) = 3.14, *p* < .001) in male participants. Overall, male participants perceived sad whole-body expressions most frequently, regardless of stimulus gender.

Likewise, female participants demonstrated similar tendencies as their male counterparts. Specifically, females reported significantly higher visibility for female sad (*M* = .44, *SEM* = .03) than female fearful (*M* = .36, *SEM* = .03, *t(*45) = −6.18, *p* < .001) or female threatening whole-body postures (*M* = .41, *SEM* = .03, *t(*45) = −2.36, *p* = .05). However, it is important to note that the latter effect was only marginally significant. In addition, visibility was significantly higher for female threatening compared to female fearful body expressions (*t(*45) = 4.32, *p* < .001) in female participants. When the effect was explored with respect to male stimuli, female participants exhibited significantly higher visibility for male sad (*M* = .47, *SEM* = .03) compared to male threatening (*M* = .40, *SEM* = .03, *t(*45) = 4.21, *p* < .001) and male fearful body expressions (*M* = .42, *SEM* = .03, *t(*45) = 2.39, *p* = .05). However, the latter effect was only marginally significant.

The emotion × stimulus gender × participant gender interaction seems not be driven by differences in stimulus visibility across emotion categories, but by differences in stimulus visibility across stimulus gender between male and female participants (Figure 2). Male participants were more likely to perceive male threatening and fearful body postures compared to female expressions. Likewise, females better perceived male than female fearful, but not threatening whole-body expressions.

Simple effects analysis was performed on the significant stimulus orientation × stimulus gender × participant gender interaction (Figure 3). Pairwise comparisons in male participants resulted in significantly higher visibility for male (*M* = .63, *SEM* = .04) in comparison with female stimuli (*M* = .60, *SEM* = .04, *t*(65) = −2.29, *p* = .022), when stimuli were presented upright. Results are comparable for inverted stimulus presentation. Males reported significantly higher visibility for male (*M* = .38, *SEM* = .04) than female whole-body expressions (*M* = .29, *SEM* = .04, *t(*65) = −4.48, *p* < .001). Interestingly, although a similar trend was present, the stimulus position × stimulus gender interaction was not significant in female participants (χ^2^(1, *N* = 45) = .09, *p* = .763).

**Figure 3.**
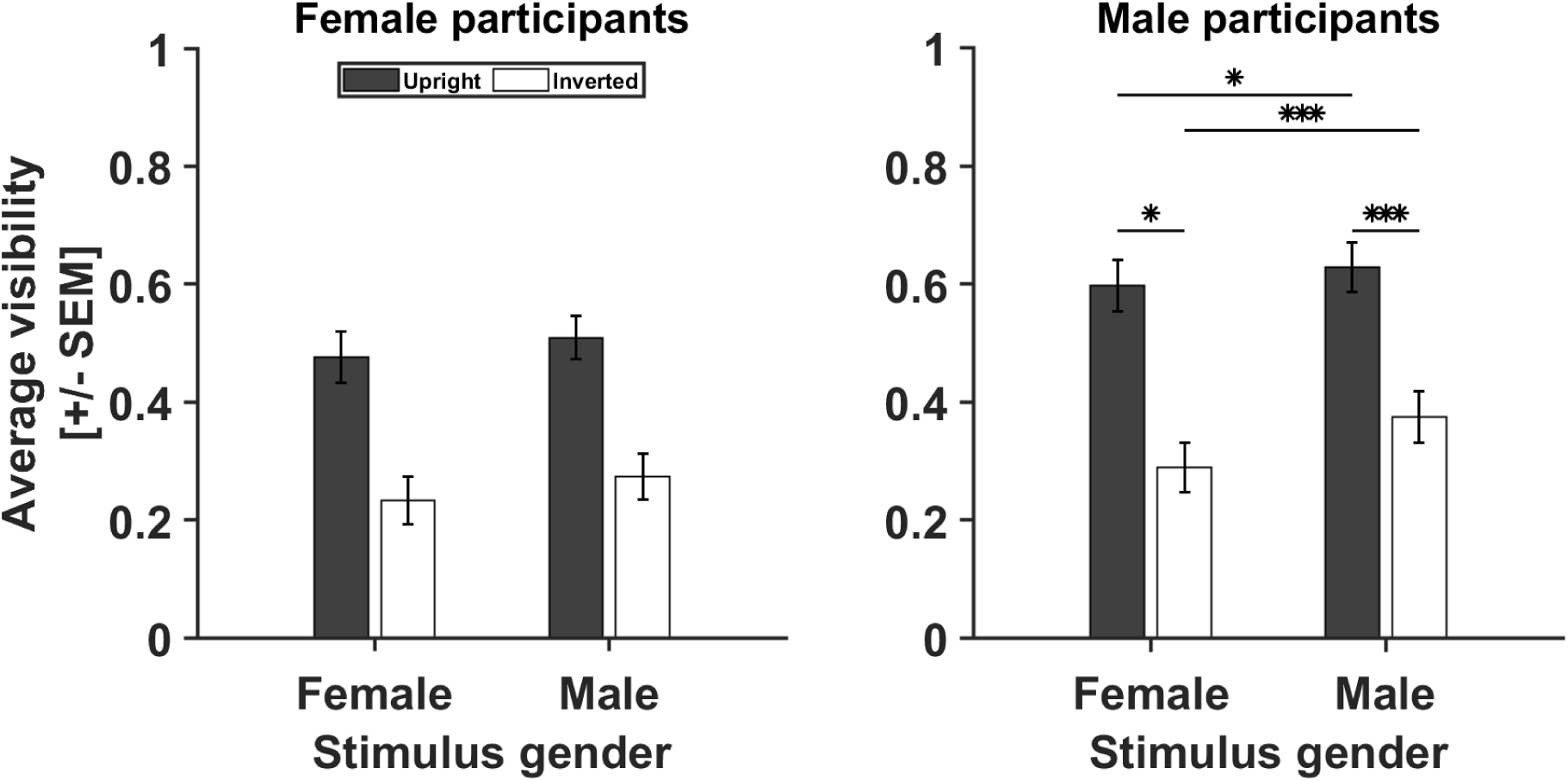
Male participants show superior visibility for male stimuli. Left and right figure represent data from male and female participants, respectively. Stimulus gender is plotted on the x-axis, whereas average stimulus visibility is plotted on the y-axis. Higher values represent higher visibility. Grey and white bars depict upright and inverted stimulus orientation, respectively. Male participants demonstrated higher visibility for male compared to female stimuli for both, upright and inverted, stimulus orientations. *SEM* = standard error of the mean, *** *p* ≤ .001, * *p* ≤ .05

In conclusion, under conditions of limited visibility male participants performed significantly better for male stimuli, independent of stimulus orientation (Figure 3). Despite displaying similar tendencies, the effect was not significant in females.

### 3.2 Reaction times

Participants’ reaction times for whole-body emotional expressions were analyzed across emotional categories, stimulus orientations as well as stimulus and participant genders. Significant main effects of emotion (χ^2^(2, *N* = 45) = 16.97, *p* < .001) and stimulus orientation (χ^2^(1, *N* = 45) = 21.15, *p* < .001) were found, as well as significant emotion × stimulus orientation (χ^2^(2, *N* = 45) = 10.45, *p* = .005), emotion × stimulus gender (χ^2^(2, *N* = 45) = 7.80, *p* < .020), emotion × participant gender (χ^2^(2, *N* = 45) = 7.01, *p* = .030), stimulus orientation × stimulus gender (χ^2^(1, *N* = 45) = 9.16, *p* = .002), stimulus gender × participant gender (χ^2^(1, *N* = 45) = 7.96, *p* = .005) and emotion × stimulus orientation × stimulus gender interactions (χ^2^(2, *N* = 45) = 6.85, *p* = .003).

On the contrary, effects of stimulus gender (χ^2^(1, *N* = 45) = 0.35, *p* = .56), participant gender (χ^2^(1, *N* = 45) = 0.75, *p* = .387) and stimulus orientation × participant gender (χ^2^(1, *N* = 45) = 2.58, *p* = .108) were non-significant.

Post hoc analysis of the emotion × participant gender interaction showed significant differences in reaction times across the three emotions in male and female participants (Figure 4). Specifically, female participants exhibited significantly faster reaction times for sad (*M* = 0.77, *SEM* = 0.05) compared to fearful expressions (*M* = 0.79, *SEM* = 0.05, *t(*91) = - 2.53, *p* = .034). Reaction time differences between sad and threatening (*M* = 0.79, *SEM* = 0.05, *t(*91) = 2.02, *p* = .126) and fearful and threatening stimuli (*t(*91) = 0.07, *p* = 1.000) were not significant.

**Figure 4.**
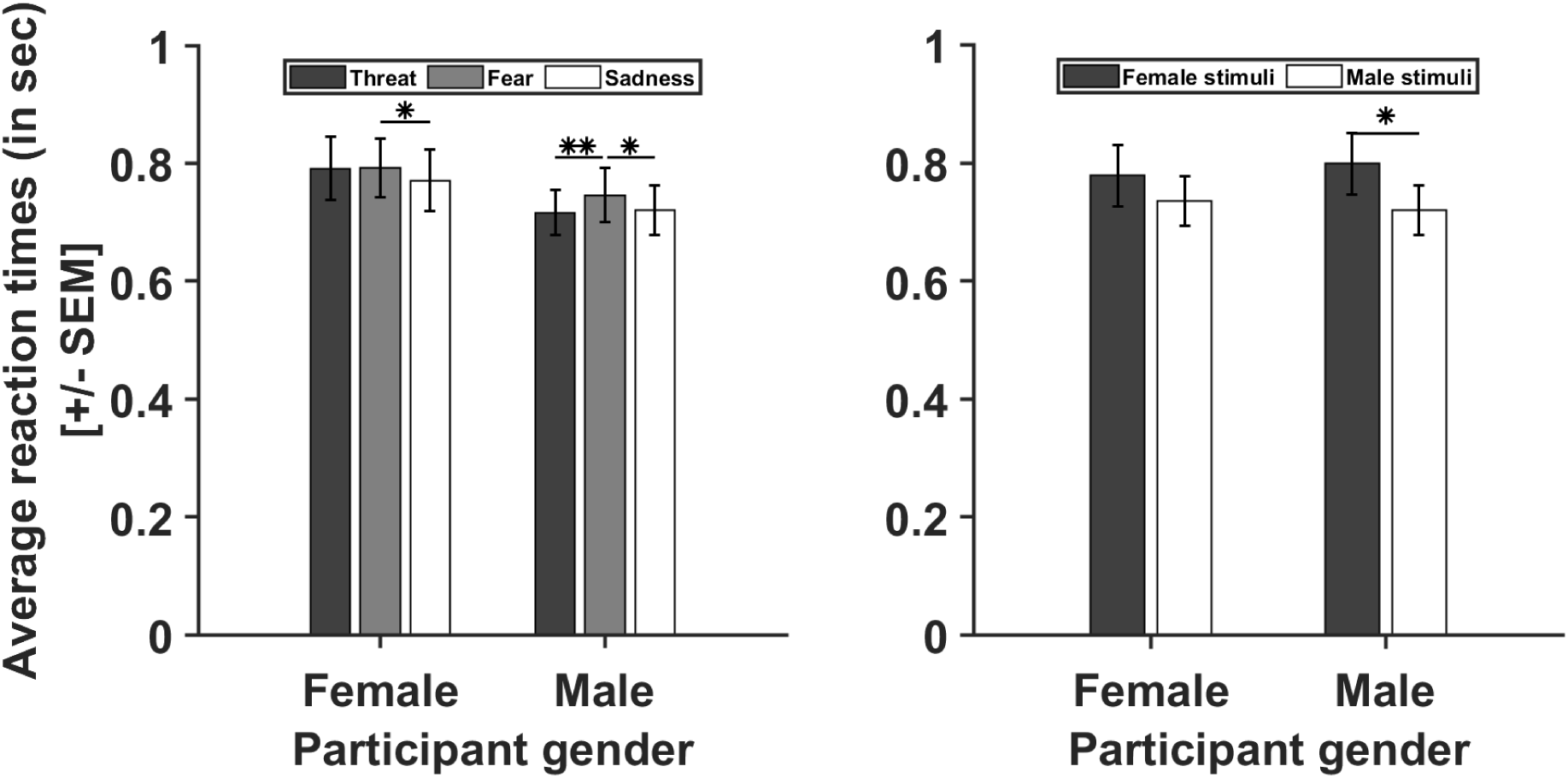
Males see other males faster. Participant gender is plotted on the x-axis, whereas average reaction times in seconds are plotted on the y-axis. **(Left)** Dark grey, light grey and white bars depict threatening, fearful and sad whole body-expressions, respectively. Female participants responded significantly faster to sad body postures than fearful ones. In male participants, reaction times were significantly faster for threatening and sad compared to fearful whole-body expressions. **(Right)** Male participants responded significantly faster to male compared to female whole-body expressions. Despite the similar trend, the effect did not reach significance in female participants*. SEM* = standard error of the mean, *** *p* ≤ .001, ** *p* ≤ .01, * *p* ≤ .05

Conversely, male participants had significantly slower reaction times for fearful (*M* = 0.75, *SEM* = 0.046) in comparison with threatening (*M* = 0.72, *SEM* = 0.04, *t(*87) = −3.27, *p* = .003) and sad body postures (*M* = 0.72, *SEM* = 0.04, *t*(87) = 2.94, *p* = .010, Figure 4). In contrast, reaction times between threatening and sad stimuli did not differ significantly (*t(*87) = 0.43, *p* = .963). Likewise, no significant differences in reaction times were observed for threatening (*t(*178) = 1.13, *p* = .257), fearful (*t(*178) = 0.66, *p* = .507) or sad whole-body expressions (*t(*178) = 0.77, *p* = .441) between male and female participants.

Overall, all participants showed faster reaction times for sad whole-body expressions under conditions of reduced visibility (Figure 4). However, unlike female participants, males also responded faster to threatening compared to fearful body postures.

We also inspected the significant stimulus gender × participant gender interaction (Figure 4). Male participants exhibited significantly faster reaction times for male (*M* = 0.72, *SEM* = 0.04) than female whole-body expressions (*M* = 0.74, *SEM* = 0.04, *t(*131) = 2.44, *p* = .015). Interestingly, female participants showed a similar trend (Figure 4). However, differences in reaction times between male (*M* = 0.79, *SEM* = 0.05) and female stimuli (*M* = 0.78, *SEM* = 0.05, *t(*137) = 1.56, *p* = .120) were not significant in female participants.

Furthermore, we conducted simple effects analysis for the significant emotion × stimulus orientation × stimulus gender interaction. For upright stimulus presentation, we observed significantly faster reaction times for male (*M* = 0.73, *SEM* = 0.03) compared to female threatening body postures (*M* = 0.76, *SEM* = 0.03, *t(*44) = 2.36, *p* = .018). Likewise, reaction times were significantly faster for male (*M* = 0.70, *SEM* = 0.03) in contrast with female sad body expressions (*M* = 0.73, *SEM* = 0.03, *t(*44) = 4.10, *p* < .001). When stimuli were inverted, the stimulus gender effect was reversed and significant only for threatening stimuli, including a similar non-significant trend for fearful stimuli. Specifically, reaction times were significantly faster for female (*M* = 0.75, *SEM* = 0.04) compared to male threatening body expressions (*M* = 0.79, *SEM* = 0.04, *t(*44) = 2.36, *p* = .020). Reaction times did not differ significantly between male and female stimuli for fearful (female: *M* = 0.76, *SEM* = 0.03, male: *M* = 0.78, *SEM* = 0.03, *t(*44) = 1.11, *p* = .268) and sad body postures (female: *M* = 0.78, *SE*M = 0.04, male: *M* = 0.77, *SEM* = 0.04, *t(*44) = 1.02, *p* = .306).

Additionally, we compared reaction times for upright and inverted stimuli across emotions. Although no significant stimulus orientation effects were observed for female threatening stimuli (*t(*44) = 0.12, *p* = .906), significantly longer reaction times were observed for inverted than upright male stimuli (*t(*44) = 3.55, *p* < .001). Likewise, reaction times significantly increased for inverted compared to upright sad body expressions for both male (*t(*44) = 4.14, *p* < .001) and female stimuli (*t(*44) = 3.13, *p* = .002). Interestingly, no significant differences in reaction times were detected between the two stimulus orientations for male (*t(*44) = 0.21, *p* = .833) or female fearful body expressions (*t(*44) = 0.65, *p* = .518).

Comparing reaction times between emotions across stimulus gender and orientation we found no significant differences in reaction times between emotions for female (χ^2^(2, *N* = 45) = 5.20, *p* = .074) and male (χ^2^(2, *N* = 45) = 1.16, *p* = .561) inverted stimuli. Nevertheless, reaction times differed significantly across emotions for male and female stimuli when stimuli were presented upright. Specifically, reaction times were significantly higher for fearful compared to sad female body expressions (*t(*44) = 2.52, *p* = .034). Otherwise, reaction times did not differ significantly between sad and threatening (*t(*44) = 1.74, *p* = .227) and fearful and threatening female stimuli (*t(*44) = 1.10, *p* = .613). In addition, reaction times were significantly higher for upright fearful compared to sad (*t(*44) = 5.13, *p* < .001) and threatening male whole-body expressions (*t(*44) = 3.11, *p* = .006). Lastly, reaction times did not differ significantly between upright sad and threatening male stimuli (*t(*44) = 2.09, *p* = .105).

## 4 Discussion

The present study investigated how specific stimulus characteristics, such as emotional expression and gender along with the gender of the participants, impact awareness. Our results indicate that participants’ awareness reflects variations in the emotion expressed, that these variations are contingent upon the gender of the stimulus but also of the participants and, finally, that minimal awareness is likely to be associated with emotion-specific stimulus features.

### 4.1 Affective and feature-based determinants of awareness

Our study uncovered significant differences between emotion categories in reaching awareness (Figure 1). Specifically, threatening expressions had an advantage over fearful ones in reaching awareness. This aligns with previous research using CFS showing that breaking from suppression is faster for threatening compared to fearful and neutral body expressions (Poyo Solanas et al., 2022a; Zhan et al., 2015). Interestingly, Poyo Solanas et al. (2023) also reported higher recognition sensitivity for fearful bodies than angry ones. One plausible explanation for these findings is that the perception of fearful body expressions might require more processing resources than expressions of threat due to their inherent ambiguity (Poyo Solanas et al., 2023). Threat signals are clear and direct, while in the case of fear expressions the locus of danger is largely unknown (Poyo Solanas et al., 2023).

Alternatively, fearful body postures may be less likely to reach awareness because they engage more direct, less cognitively demanding subcortical pathways. Indeed, increased functional connectivity between the right amygdala, pulvinar and superior colliculus has been reported for ‘unseen’ compared to ‘seen’ fear-conditioned faces, whereas right amygdala connectivity with the fusiform and orbitofrontal cortex decreased for ‘unseen’ stimuli (Morris, Öhman and Dolan, 1999). Numerous other studies in neurotypicals (Tamietto and de Gelder, 2008) and blindsight patients (de Gelder, Morris and Dolan, 2005) have shown that fearful facial expressions can be processed without visual awareness. Likewise, fearful body expressions can be non-consciously processed in blindsight (Tamietto et al., 2009) and intact brains (Zhan and de Gelder, 2019). This suggests that subcortical processing of fearful body expressions may take precedence for adaptive behavior, possibly at the expense of triggering stimulus awareness, which typically involves cortical structures. In further support of this notion, we observed longer reaction times for fearful whole-body expressions compared to other emotion categories in male participants (Figure 4), indicating possible freezing behavior. A related study investigating facial expressions found that recognition of fearful faces was the least accurate and the slowest among seven emotion categories (Calvo and Lundqvist, 2008). Similarly, Poyo Solanas et al. (2022) associated freezing behavior with slower heart rate when participants were exposed to fearful compared to neutral body expressions in conditions of reduced visual awareness. Moreover, several studies reported suppression of motor readiness following exposure to fearful body expressions (Borgomaneri et al., 2015; Borgomaneri, Vitale and Avenanti, 2017). Taken together, these findings suggest that fearful body expressions do not necessarily have to become fully aware to be processed. Nonetheless, female participants’ reaction times were comparable for threatening and fearful body expressions (Figure 4). Therefore, further research is necessary to shed additional light on these findings and potential gender differences in freezing behavior when confronted with threatening and fearful social signals.

Surprisingly, sad body postures accessed visual awareness more frequently (Figure 1) and were accompanied with faster reaction times than threatening and fearful expressions (Figure 4). Considering that sadness has been rarely investigated in previous studies, little is known about the perception of sad body expressions under minimal awareness. One possible explanation is that sadness fails to trigger an action plan due to a lack of danger signaling, whereas threat and fear require behavioral reactions to increase survival. Indeed, it has been shown that fearful whole-body expressions induce fear contagion and increased activation in action preparation areas, i.e., the supplementary motor area (SMA) and precentral gyrus (de Gelder et al., 2004). Likewise, a virtual reality study investigating freezing behavior under threat reported a decrease in heart rate when participants were armed and able to shoot an opponent in comparison to being unarmed in a virtual scenario, suggesting that freezing behavior could be related to action preparation (Gladwin et al., 2016). Therefore, it is possible that, unlike threatening and fearful, sad body expressions are less likely to trigger defensive motor preparation, resulting in faster reaction times for sad stimuli.

Lastly, to advance our currently limited understanding of the detailed visual processes involved in non-conscious perception, we contrasted performance for upright opposed to inverted stimuli to explore the contribution of configural vs. feature-based processing in minimal awareness. Consistent with Stein et al. (2012), we found a substantial awareness reduction for inverted presentation (Figure 1) across the three expression categories. This may indicate that lower stages of awareness are associated with feature-based processing, as the differences between the expressions are expression-defining features. Namely, postural features, such as limb contraction, can distinguish between various whole-body expressions and drive perception (Poyo Solanas et al., 2020a,b). Likewise, it has been argued that feature-based perception may be sufficient for non-conscious or minimally conscious affective perception (de Gelder and Poyo Solanas, 2021). However, whether the observed performance drop for inverted stimuli is due to a loss of configural processing associated with unawareness/minimal awareness or feature based processing only is a matter for future research.

### 4.2 Gender specificity

Our results clearly show a gender effect on awareness of the stimuli as well as of the participants. In male participants, male whole-body expressions reached visual awareness more often compared to female stimuli, independent of stimulus orientation (Figure 3) and this difference was not significant in female participants (Figure 3). Reaction times are consistent with this gender specific pattern, with male participants having faster reaction times for male body postures in comparison with female stimuli, whereas there was a similar but non-significant trend in female participants (Figure 4).

Regarding specific emotion effects, male participants were more susceptible to male expressions of threat and fear (Figure 2). In contrast, females were more likely to perceive male expressions of fear only (Figure 2). These findings are likely a consequence of gender differences in neural processing of threatening body signals. In this regard, Kret et al. (2011) found higher BOLD responses in the STS, EBA and pre-SMA of male participants, when male participants were observing male dynamic whole-body expressions of threat. Unlike females, male participants also showed a motor preparation response for threatening male body postures in the premotor cortex (Kret et al., 2011). Interestingly, research on faces showed that, in general, male angry facial expressions are better recognized than female (Calvo and Lundqvist, 2008).

Taken together, these observations indicate an evolutionary relevance of male threatening signals, especially to males. Tay (2015) argued that, from an evolutionary point of view, male and female emotional facial expressions have different adaptive values. Thus, perceiving male expressions of threat and, even more so, perceiving them rapidly may benefit survival. This may also explain female sensitivity towards fearful male body expressions and faster reaction times for male stimuli in both genders, albeit non-significantly in females (Figure 4).

Additionally, men show greater sensitivity to threatening cues in general and this may be mediated by testosterone levels (Kret and de Gelder, 2012). Testosterone levels correlate positively with activity in the amgydala and orbitofrontal cortex and are positively associated with alertness towards angry facial expressions in males and females (Kret and de Gelder, 2012). Likewise, testosterone has been shown to enhance the effects of vasopressin, a hormone regulating the fight-or-flight response, which thereby shapes defensive behavior in male animals (Kret and de Gelder, 2012). In rodents, estrogen and testosterone mediate the function of oxytocin (OT) and arginine vasopressin (AVP) in social recognition of conspecifics (Aspesi and Choleris, 2022). Estrogen has facilitatory effects on social recognition of male and female rodents by regulating OT action in the hypothalamus and medial amygdala, whereas the AVP system comprises the stria terminalis, amygdala and the lateral septum and plays a more important role in social recognition of male rodents (Aspesi and Choleris, 2022). OT levels are influenced by estrogen and it is believed to be involved in attenuating stress, while AVP seems to be strongly dependent on testosterone levels and has anxiety-inducing effects (Bos et al., 2012). Interestingly, AVP release in the septum seems to facilitate simple stimulus–response associations, while interfering with complex stimulus processing, indicating that AVP promotes goal-directed behavior supporting more effective responses to threat (Bos et al., 2012, Engelmann, 2008, Engelmann and Landgraf, 1994). Taken together, this complex interplay of testosterone and AVP levels could possibly explain why males respond faster to male whole-body emotional expressions. Nonetheless, further research is necessary to shed light on the complex function of sex steroids in regulating brain areas involved in emotional processing.

In conclusion, our results indicate that awareness needs to be understood in relation to the specific affective expression of the stimuli, to the gender as well as to the gender of the participants. We found that threatening whole-body expressions reach visual awareness more easily than fearful body postures, and that males displayed higher susceptibility for male body expressions of threat and fear. Our results provide novel insights on the role of gender in emotional processing, highlighting its clinical importance for standard investigations of face and body emotion recognition. We suggest that the present findings about affective and gender determinants of awareness are compatible with theories that underscore biological and somatic determinants of awareness (Panksepp, 2007; Damasio, 1999). Based on these theories, one expects that subjective awareness reflects biological and somatic characteristics of perceivers.

## 5 Limitations

While the current study offers valuable insights into the intricate interplay of emotional visual processing, gender cues, and individual characteristics, it is important to acknowledge certain limitations inherent in the research design. One notable limitation lies in the assessment method of subjective awareness. We chose a binary outcome measure (seen/unseen) which might have oversimplified the nuanced nature of perception. Future studies should consider incorporating finer measures that may better capture subjective perceptual experience and the possible different degrees of perceptual awareness. In addition, the absence of conditions contrasting whole bodies versus body parts might have hindered a comprehensive exploration of configural versus part-based processing.

## 7 Author Contributions

EJ: Conceptualization, Methodology, Data collection, Data analysis, Writing—original draft, review & editing, MPS: Conceptualization, Methodology, Data analysis, Writing—review & editing), BdG: Conceptualization, Funding acquisition, Project administration, Supervision, Writing—original draft, Writing—review & editing.

## 9 Acknowledgments

The authors thank Borgomaneri et al. (2023) for their help with task development.

